# LC/MS-QToF Profiling, Anti-Diabetic and Anti-Adipogenic potential of Divya MadhuKalp: A Novel Herbo-mineral Formulation

**DOI:** 10.1101/2020.02.15.950865

**Authors:** Acharya Balkrishna, Alpana Joshi, Subrata K. Das, Laxmi Bisht, Sachin Sakat, Vinamra Sharma, Niti Sharma, Khemraj Joshi, Sudeep Verma, Vinay K Sharma, CS Joshi

## Abstract

The incidence rate of diabetes mellitus is increasing worldwide. Herbal formulations have recently gained importance as an alternative therapeutic option in controlling diabetes without causing any side effects. In the present study, we have demonstrated maintenance of glycemic homeostasis and anti-adipogenic potential of a herbo-mineral formulation Divya MadhuKalp (DMK). Initially, we evaluated the presence of bioactive compounds in DMK using LC/MS-QToF analysis. In-vitro analysis of DMK in L6 (skeletal muscle) cells showed a significant increase in cellular glucose uptake. Similarly, a human equivalent dose of DMK significantly reduced blood glucose level in normoglycemic and oral glucose tolerance rat model. DMK extract also inhibited formation of advanced glycation end product and showed anti-α-glucosidase activity. Further analysis of DMK in 3T3 L1 pre-adipocytes demonstrated anti-adipogenic activity through reduction in intracellular lipid accumulation and triglyceride contents along with downregulation of major adipogenic transcriptional factors (PPAR-γ and C/EBPα) and, adipocytes marker genes (LPL, AP2 and adiponectin). In conclusion, DMK exhibited anti-diabetic and anti-adipogenic activities by synergistic effect of its bioactive compounds and can be considered as a potent herbo-mineral formulation for treating metabolic diseases.

## Introduction

Diabetes mellitus (DM) is a chronic metabolic disorder that has profound effects on patient’s quality of life in terms of health and socio–economic levels. According to International Diabetes Federation’s (IDF) projections, approximately 693 million people will be diagnosed with DM by year 2045 ^1^. DM is characterized by abnormally high blood glucose resulting from defective metabolism of carbohydrates, lipids and proteins. Long-term pathophysiological complications associated with DM are cardiovascular diseases, nephropathy, neuropathy and retinopathy. During hyperglycemic condition, there is also development of advanced glycation-end products (AGEs) as a consequence of protein glycation. AGEs are the possible causal factor lead to the development of diabetes related complications. Currently, the core objective of diabetes treatment is to maintain the normal blood glucose levels and, to prevent or delay its metabolic complications.

Obesity is the excessive accumulation of fat or adipose tissue in the body ^2^. It is a major risk factor for the development of type 2 diabetes mellitus (DM). Adipogenesis involves differentiation of pre-adipocytes into mature adipocytes and is regulated by two major adipogenic transcription factors namely, Peroxisome proliferator–activated receptor **γ** (PPARγ) and CCAAT/enhancer binding protein **α** (C/EBPα) and downstream induction of adipocyte-related genes including, adipocyte protein 2 (AP2), lipoprotein lipase (LPL), adiponectin etc.^3, 4, 5^.

Medicinal plants have great importance to human health and communities. The medicinal value of these plants typically results from the presence of many bioactive compounds. In contrast to synthetic drugs based on single molecule, many bioactive compounds of poly herbal formulation exert their beneficial effects through the additive or synergistic action acting at single or multiple target sites associated with a physiological process^6, 7^. Unlike, synthetic drugs, which may induce unwanted side-effects, natural herbal formulations tend to show similar efficacy with minimal to no side-effects and cost effectiveness^6, 7, 8^. In traditional Indian medicine system, numerous herbal and herbo-mineral formulations have been used for treating various diseases, but their scientific evidence is still lacking. Divya MadhuKalp (DMK) is an Ayurvedic herbo-mineral formulation has been extensively used in clinical practices for managing blood glucose levels during onset of DM. The DMK comprises of eight herbs and one minerals–rich organic component in specific proportion to make complex bioactive formulation (Table 1). The ingredients are; *Momordica charantia* ^9^; *Picrorhiza kurroa*^10^; *Swertia* chiraita ^11^; *Azadirachta indica* ^12^; *Trigonella foenum-graecum* ^13^; *Syzyium cuminii* ^14^; *Withania somnifera* ^15^; *Aconitum hetrophyllum* ^16^ and *Shilajeet* (Asphaltum) ^17, 18^. The ingredients in DMK have been used for diabetic therapy in ancient Indian Ayurvedic medicinal system. In order to explore the pharmacological mechanism of poly herbal extracts, accurate characterization of the bioactive compounds is very essential. Moreover, the quality control is very important factor in discovery of a new drug. LC/MS-QToF technology is most appropriate analytical method for the full characterization and quality control of poly herbal extract ^19^. DMK associated herbal ingredients have been found to be rich in secondary metabolite compositions such as, phenolics, terpenes, and nitrogen containing compounds. These active bio-molecules play an important role in carbohydrate and lipid metabolism leading to efficient management of diabetes progression and associated complications ^20, 21^.

**Table 1.**
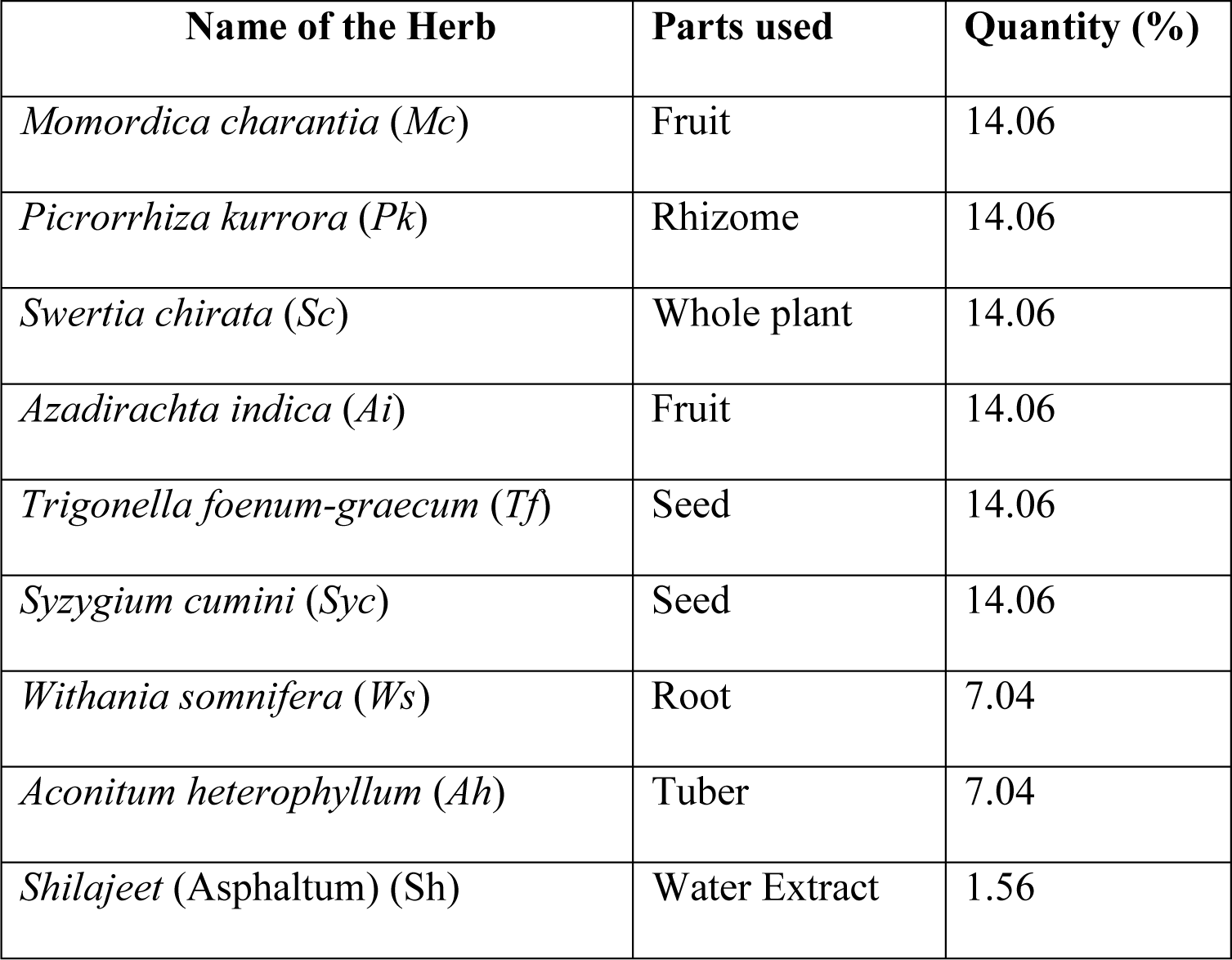
Composition of Divya MadhuKalp (DMK)

In the present study, we performed the chemical profiling by LC/MS-QToF analysis and determined the anti-diabetic and anti-adipogenic potential of the herbo–mineral extract of DMK using *in vitro* and *in vivo* models.

## Material and Methods

### Experimental Material

Divya MadhuKalp (DMK) was sourced from Divya Pharmacy, Haridwar, India. Dulbecco’s Modified Eagle Medium (DMEM), fetal bovine serum (FBS), Insulin and bovine serum albumin (BSA), MTT [3-(4, 5-dimethylthiazol-2-yl)-2, 5-diphenyltetrazolium bromide], IBMX (3-isobutyl-1-methylxanthine), dexamethasone were obtained from (ThermoFischer, USA). 2-(N-(7-Nitrobenz-2-oxa-1,3-diazol-4-yl)amino)-2-deoxy-d-glucose (2-NBDG) was purchased from Invitrogen (Carlsbad, CA, USA). Trypsin-EDTA solution was purchased from GIBCO BRL (Grand Island, NY, USA). Sterile 96 well plates were obtained from Tarsons India Pvt. Ltd. (Kolkata, India). Oil-Red-O, metformin, α-glucosidase, acarbose, p-nitrophenyl glucopyranoside, aminoguanidine, glibenclamide, D-glucose, and sodium carboxy methyl cellulose were purchased from Sigma-Aldrich (USA). The glucose monitoring system, measurements strips (Life Scan, Switzerland) and sodium chloride injection I.P. 0.9 % w/v (Infutec Healthcare Ltd. India) was procured locally. All other chemicals and reagents used were of the highest commercial grade. HPLC grade reagents of analytical grade toluene, ethyl acetate, formic acid, Gallic acid Vanillic Acid, Caffeic acid, Trigonelline, Adinosine, Rutin Quercetin Picriside I Pcroside II, Trans-Cinnamic acid and Apigenin were obtained from the Sigma-Aldrich, India.

### Experimental Animals and Reagents

Male Wistar rats (160-180 g) were procured from Liveon Biolabs Pvt. Ltd., Bengaluru, India. Animals were housed in polypropylene cages in controlled room temperature 22 ± 1 °C and, relative humidity of 60-70 % with 12:12 h light and dark cycle in a registered animal house (1964/PO/RC/S/17/CPCSEA). The animals were supplied with standard pellet diet (Golden Feed, New Delhi, India) and sterile water *ad libitum*. The animal experimental study protocol for this work was approved by the Institutional Animal Ethical Committee of Patanjali Research Institute, India (IAEC protocol number- PRIAS/LAF/IAEC-003) and all the experiments were performed following relevant guidelines and regulations.

### Preparation of Divya MadhuKalp (DMK) Extract

DMK (20 g) and its individual constituents were suspended separately in 300 ml of 70 % aqueous ethanol. The suspension was refluxed for 6 h at 45 °C and, filtered through Whatman filter no-41. The filtrates were evaporated at 45 °C under reduced pressure in a rotary evaporator. The final dried samples were stored in desiccator at room temperature.

### High-Performance Liquid Chromatography (HPLC) Analysis

The extract of DMK was diluted in 50 % methanol (1 mg/ml) and subjected to HPLC analysis. Waters binary HPLC system (Waters Corporation, Milford, MA, USA), equipped with, column oven, auto–sampler (Waters 2707) and, photodiode array (PDA) detector (Waters 2998) was used for the analyses. A reversed phase C18 analytical column (4.60×250 mm, 5 μm particle size; Sunfire, Waters, USA) was utilized at 30 °C column temperature. The injection volume was 10 µl (DMK) and 10 µl (standard mix) at different concentration. The chromatograms were acquired at 230 nm according to the absorption maxima of the analyzed compounds. Each compound was identified by its retention time (RT) and, by spiking with standards under the same conditions. The identities of constituents were also confirmed with a photodiode array (PDA) detector by comparison with ultraviolet (UV) spectra of standards in the wavelength range of 210–650 nm.

### Liquid Chromatography Mass Spectroscopy-Quadrupole Time-of-Flight (LC/MS-QToF) Analysis

Twenty milliliter of methanol: water (60:40) was added in 200.3 mg of powdered Divya MadhuKalp (DMK) and sonicated for 30 min. Then this solution was centrifuged for 5 min at 10,000 rpm and filtered through 0.22 µm nylon filter. The extract was used for LC/MS analysis The LC/MS-QToF instrument was equipped with an ESI ion source operating in a positive and negative ion mode. A mass range of 50-1000 Da was set with a 0.2 s scan time. The main working parameters for mass spectrometry were set as follows, ionization type-ESI, mode-MS^E^, acquisition time-45 min, mass range (*m*/*z*) 50–1000 *m*/*z*, low collision energy-6 eV, high collision energy 20-40 eV (ramp), cone voltage-40 V, capillary voltage-1 kV (for positive ion mode) (Figure 1A), capillary voltage-2 kV (for negative ion mode) (Figure 1B), source temperature-120 °C, desolvation temperature-450 °C, cone gas flow-50 L/h, desolvation gas flow-900 L/h. Mass was corrected during acquisition, using an external reference (Lock-Spray) consisting of 0.2 ng/ml solution of leucine enkephalin infused at a flow rate of 10 μl/min via a lock-spray interface, generating a reference ion for the positive ion mode [(M + H^+^) + *m*/*z*556.2766] and for the negative ion mode [(M - H^+^) *m*/*z* 554.2620] to ensure mass accuracy during the MS analysis. The Lock-Spray scan time was set at 0.25 s with an interval of 30 s. Analysis was performed on a Waters Xevo G2-XS QToF equipped with Acquity UPLC-I Class and Unifi software. Separation was carried out using Acquity UPLC HSS-T3 column (100 × 2.1 mm, 1.7 µm). The column was maintained at 40 °C throughout the analysis, and sample temperature was kept at 20°C. The elution was carried out at a flow rate of 0.4 ml/min using gradient elution 0.1 % formic acid in water (mobile phase A) and 0.1 % formic acid in acetonitrile (mobile phase B). Solvent gradient program was 95 %-90 % of the mobile phase A during 0-5 min, 90 %–80 % A during 5-10 min, 80 %-60 % A during 10-20 min, 60 %-40 % A during 20-30 min, 40 % A during 30-45 min, 40 %-95 % A during 45-46 min, followed by 95 % A during 46-50 min. A total of 2 µl of the test solution was injected for the screening and the chromatograph was recorded for 45 min.

**Fig 1.**
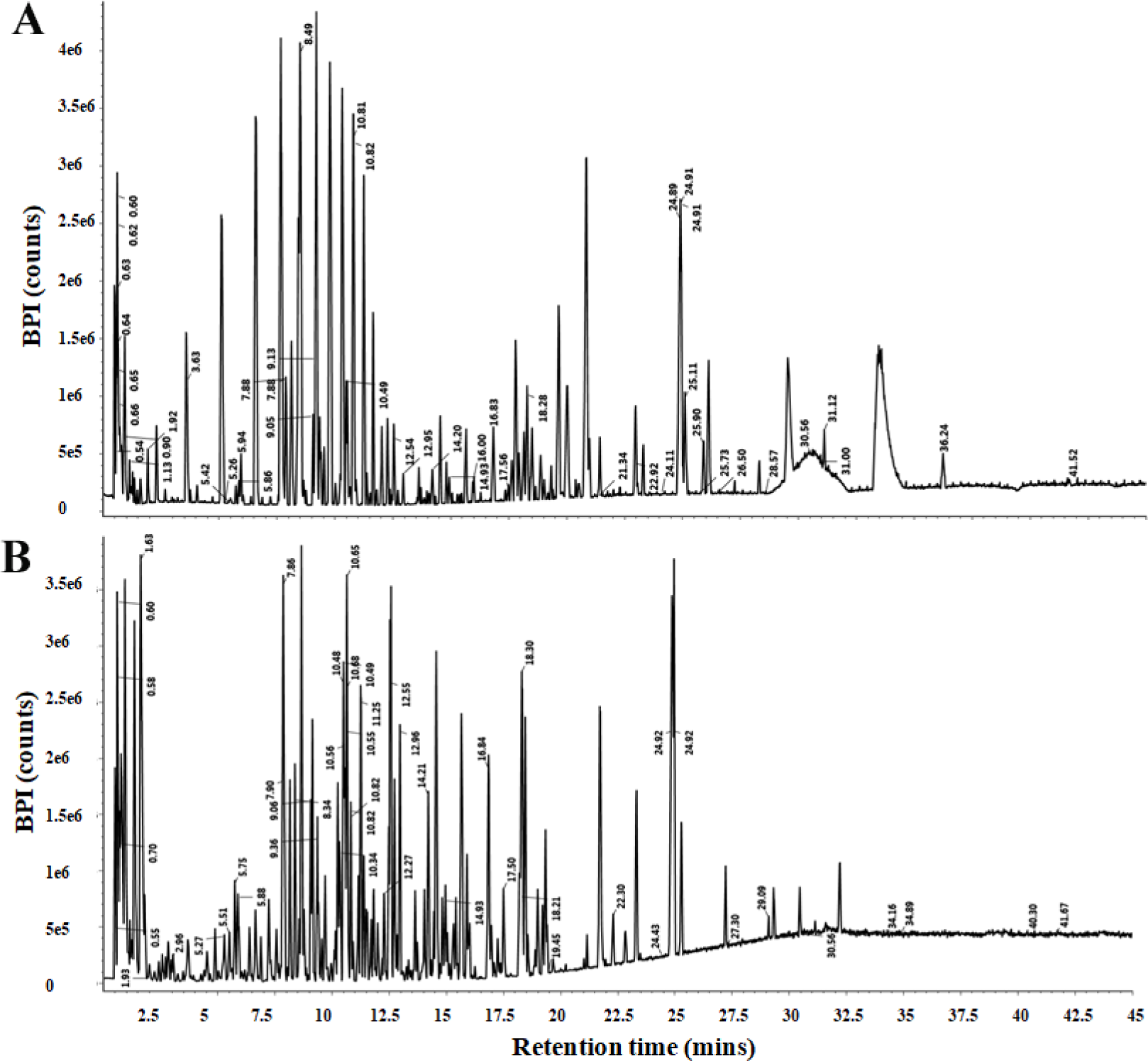
Base peak chromatogram of hydroalcoholic extract of Divya MadhuKalp (DMK). (A) Positive and (B) Negative ion modes by LC/MS-QToF. The labels of the total compound chromatogram peaks are corresponding to the compound retention time in Table 2

**Table 2.**
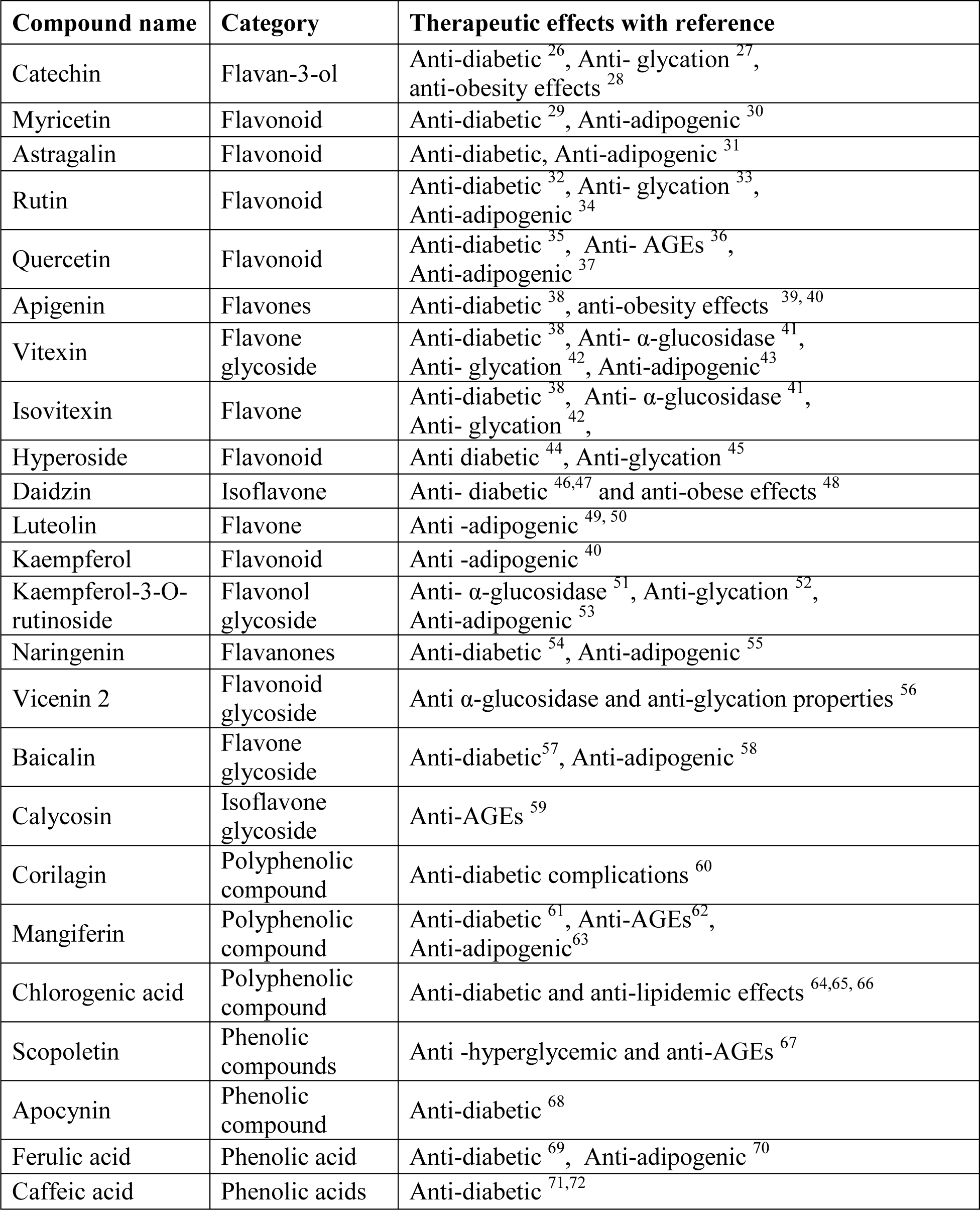
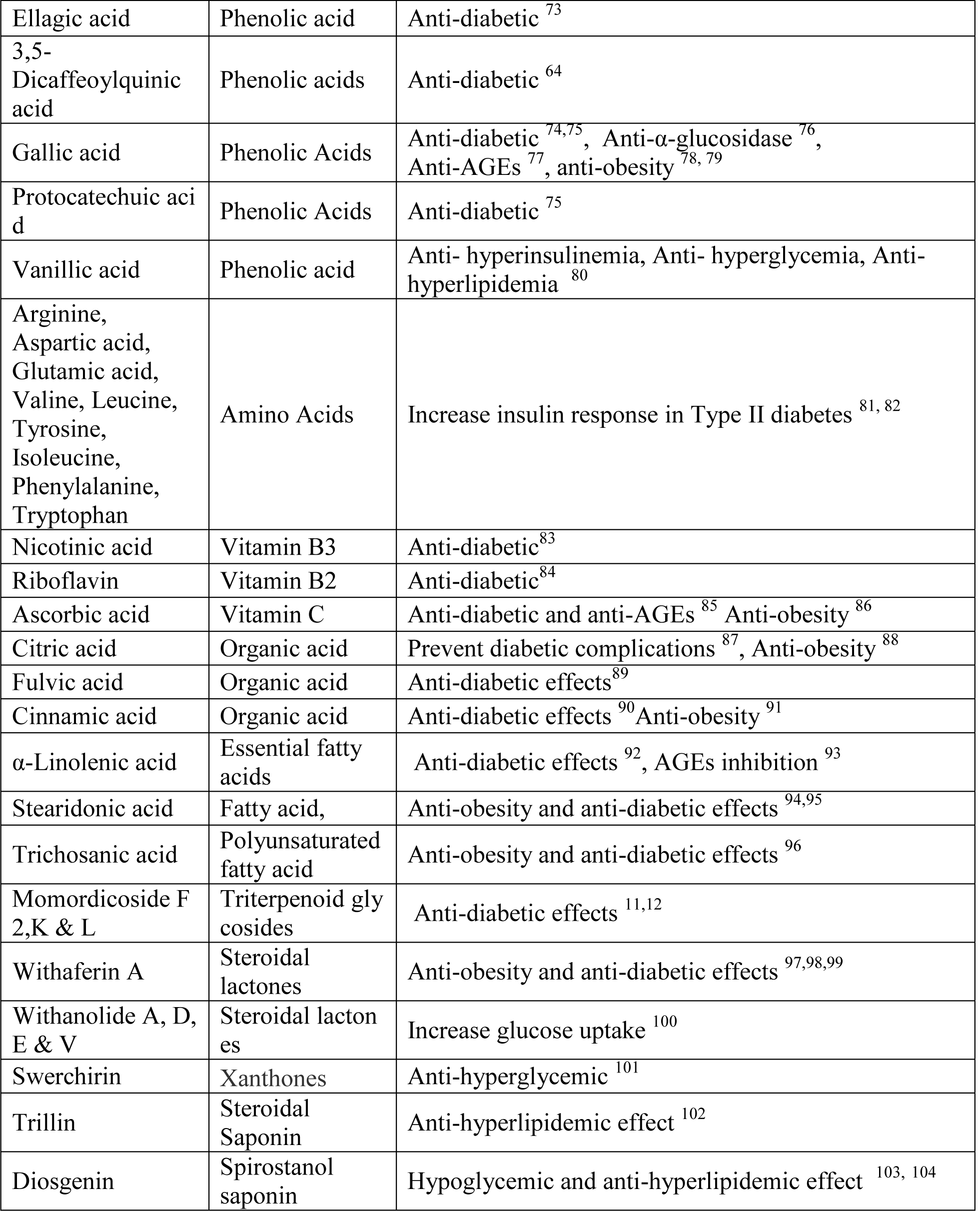
Phytochemical category and therapeutic effects of referenced bioactive compounds present in Divya MadhuKalp (DMK)

### Cell lines and Culture condition

Mouse 3T3-L1 pre-adipocytes and rat skeletal muscle L6 cell lines were procured from ATCC licensed repository, National Centre for Cell Sciences (NCCS), Pune, India. Cells were cultured in 96 well culture plates with complete medium (high glucose Dulbecco’s Modified Eagle Medium, 10 % fetal bovine serum (FBS) and 50 μg/ml penicillin/streptomycin).

### In–vitro Cytotoxicity Assay

Cells (3T3-L1 and L6) were seeded in a 96-well plate (5 × 10^3^ cells/well) and incubated at 37 °C, 5 % CO_2_ with complete medium for 24 h. After 24 h, the medium was changed to complete medium supplemented with DMK extract and its constituents at various concentrations for 24 h. MTT solution (5 mg/ml in PBS) was then added to the plate (10 μl/100 μl medium/well) and incubated for 4 h at 37 °C. The resultant formazan product was dissolved in DMSO (200 μl/well) and absorbance was measured at 492 nm by EnVision Multimode Plate Reader (Perkin Elmer, USA).

### Glucose Uptake in L6 cells

L6 cells were cultured in a humidified atmosphere of 5 % CO_2_ at 37 °C for 2 days to grow up to 70 % confluency. The cells were dissociated with Trypsin-EDTA solution (GIBCO BRL, Grand Island, NY, USA) and, seeded in 96 micro well plates (Tarsons India Pvt. Ltd., Kolkata, India). After two days incubation, cells were starved with serum and glucose free DMEM media for 2 h. Cells were treated with different concentrations of DMK extract (0-250 µg/ml) and single concentration (62.5 µg/ml) of ten individual extracts (DMK and its nine components) in glucose free media containing 0.05 % FBS and insulin (100 nM). 2-NBDG was added (50 μg/ml) in the culture, incubated at 37 °C for 30 min. Metformin (Sigma-Aldrich, USA) was used as a positive control at the concentration of 100 µM. After incubation, cells were harvested, washed thrice with PBS and, the resultant fluorescence was measured with flow cytometer (Amnis® Imaging Flow Cytometers, Millipore, USA). Cells taking up 2-NBDG displays fluorescence with excitation and emission at 485 nm and 535 nm, respectively, and, measured in appropriate channel. For a given measurement, data from 8,000 single cell events was captured using flow cytometry.

### Alpha-Glucosidase Activity

The effect of DMK extracts on alpha-glucosidase activity was determined using alpha– glucosidase (Sigma-Aldrich, US) from *Saccharomyces cerevisiae*. Acarbose was used as a positive control. The enzyme (0.33 µg/ml) was pre–incubated with the extracts at different concentrations for 10 min. The reaction was initiated by adding 3 mM p-nitrophenyl glucopyranoside (pNPG) and the reaction mixture was incubated at 37 °C for 20 min. The reaction was terminated by adding 0.1 M Na_2_CO_3_^+^ solution (500 µl). The alpha-glucosidase activity was determined by measuring the yellow-colored para-nitrophenol released from pNPG at 405 nm (EnVision Multimode plate reader, Perkin Elmer, USA). The results were expressed as the percentage of the blank.

### Anti-Glycation Activity

The anti-glycation activity of DMK extract was determined using the bovine serum albumin (BSA) (Hi-Media, India) following the method described with slight modification ^22^. BSA glycation reaction was carried out by incubating 1 ml of 50 mg/ml BSA in 0.1 M phosphate buffer (pH 7.4) and, 0.5M dextrose monohydrate containing 5 mM sodium azide as bacteriostatic at 37 °C for 1-4 weeks with various concentrations of the extracts. Aminoguanidine (Sigma– Aldrich, USA) was used as a positive control. The BSA glycation was monitored at 370/440 nm by using EnVision Multimode Plate Reader (Perkin Elmer, USA).

### Dose Selection and Preparation of Test Formulation

The human equivalent dose of DMK for the rat experiments was estimated through body surface area ^23^. The human recommended dose of DMK is 500-1000 mg twice a day. Therefore, maximum equivalent dose for rat was estimated as 200 mg/kg/day, considering dose conversion factor of 6.2 ^23^. The powdered form of DMK (Batch no-MNVE004) was suspended in 0.25 % Na-CMC, triturated to form uniform suspension and used for the *in-vivo* experiments.

### Assessment of Hypoglycemic Activity in Normal Rats

The acute effect of single dose exposure of DMK was evaluated for hypoglycemic potential in normal animals according to the modified method ^24^. All the animals were fasted for 14-16 h before commencing the experiment. Fasting blood glucose levels (0 min) were measured using glucose testing strips and, randomized into different groups of eight animals each. Control animals were treated orally with 0.25 % Na-CMC. The standard drug Glibenclamide (GLB) or Test sample DMK was administered orally to animals at 10 or 200 mg/kg doses, respectively. Blood glucose level was measured at 30, 60, 90 and 120 min after the drug treatment using tail snip method.

### Assessment of Hypoglycemic Activity by Oral Glucose Tolerance Test (OGTT) in normal rats

Oral Glucose Tolerance Test was performed on 14-16 h fasted rats ^25^. Control, reference control and, treatment animals administered orally with 0.25 % Na-CMC, GLB (10 mg/kg) and DMK (200 mg/kg), respectively to assess the hypoglycemic activity of DMK. After 30 min of drug treatment, all the animals treated orally with D-Glucose at a dose of 3 g/kg; and blood glucose levels were measured at different time points (30, 60, 90 and 120 min).

### 3T3-L1 Adipocyte Differentiation

Mouse 3T3-L1 cells were grown in complete media and upon confluency, the media were replaced with adipogenic differentiation cocktail media (MDI) containing Insulin (10 μg/ml), 0.5 mM IBMX, 1 mM dexamethasone on the following day ^4^. At the same time, cells were treated with various concentrations of DMK extract (0-250 µg/ml) and single concentration (250 µg/ml) of ten extracts (DMK and its nine components), except *Trigonella foenum-graecum, Tf* (62.5µg/ml). The culture was further incubated at 37 °C, 5 % CO_2_ for 2 days. After that, the induction media were changed with differentiation media (DMEM + 10 % FBS + 10 μg/ml Insulin) for 6 days. At this point media was changed every alternate day. Adipocytes were treated with DMEM media (DMEM + 10 % FBS) for 2 days to mature the oil droplets. The extent of differentiation was determined on day 10 by Oil Red O staining.

### Oil Red O Staining

Oil Red O staining on day 10 after induction determined intracellular lipid accumulation. After removal of culture media, the cells were washed twice with PBS, fixed with 10 % formalin and, stained with Oil Red O (six parts 0.6 % Oil red O dye in isopropanol and four parts water) for 30 min. After three rinses with distilled water and, once with 60 % isopropanol, these cells were photographed under the bright field microscope (Primovert, Zeiss, USA). To quantify the lipid accumulation, lipid and Oil Red O were dissolved in isopropanol and, absorbance was measured by a microplate spectrophotometer at 495 nm (EnVision Multimode plate reader, Perkin Elmer, USA). The percentage of Oil red O stained material relative to control wells was calculated.

### Quantification of Triglyceride Deposition

Intracellular triglycerides (TG) content was measured using Triglyceride Colorimetric Assay Kit (Cayman chemicals, USA) according to the manufacturer’s protocol. In brief, cells were washed with PBS, harvested by trypsinization and, re–suspended in PBS. The cell suspension was homogenized by sonication, the enzymatic reaction was carried out for 15 min at room temperature and finally, the absorbance (540 nm) was taken using plate reader (EnVision Multimode plate reader, Perkin Elmer, USA).

### Quantitative Real–Time PCR Analysis

Total RNA was extracted (RNeasy Mini Kit, Qiagen, USA) from mouse 3T3-L1 pre-adipocytes at the desired 4^th^ day of adipogenic differentiation, treated with various concentrations (0-250 µg/ml) of DMK. First strand cDNA was synthesized using 1 µg of total RNA using manufacturer protocol (Verso cDNA Synthesis Kit, ThermoFischer, USA). Selected genes were amplified and, quantified by performing PCR reaction using Brilliant II SYBR® Green QPCR Master Mix (Agilent Technologies, USA). The primer sequences for PCR analysis were as follows: PPARγ (sense) TTCAGCTCTGGGATGACCTT, (antisense) CGAAGTTGGTGGGCCAGAAT; C/EBPα (sense) GTGTGCACGTCTATGCTAAACCA, (antisense) GCCGTTAGTGAAGAGTCTCAGTTTG; LPL (sense) GGCCAGATTCATCAACTGGAT, (antisense) GCTCCAAGGCTGTACCCTAAG; Adiponectin (sense) GTTGCAAGCTCTCCTGTTCC, (antisense) ATCCAACCTGCACAAGTTCC; AP2 (sense) CATCAGCGTAAATGGGGATT, (antisense) TCGACTTTCCATCCCACTTC, and, GAPDH (sense) AAGAAGGTGGTGAAGCAGGCATC, (antisense) CGAAGGTGGAAGAGTGGGAGTTG. PCR conditions were as follow: 1 cycle of 50 °C for 10 min and 95 °C for 5 min, 35 cycles of 95 °C for 10 s, 60 °C for 30 s and 72 °C for 30 s. Relative gene expression was expressed as relative mRNA level compared with the control, was calculated after normalization to GAPDH (house-keeping gene) following the 2^ΔΔCT^ method.

### Statistical Analysis

All experimental data are presented as the mean ± standard error of the mean (SEM). Statistical analysis was done using GraphPad Prism version 7.0 software (La Jolla, CA, USA). A one–way analysis of variance (ANOVA) followed by Dunnett’s multiple comparison t–test was used to calculate statistical difference. Statistical significance was considered at *p < 0.05 or **p <0.01.

## Results

### Identification of Marker Compounds of Divya MadhuKalp (DMV)

A detailed phytochemical profiling of DMK was performed using LC/MS-QToF analysis. A total 139 compounds have been identified. The chromatogram of the hydroalcoholic extract from DMK was displayed in **Figure 1** (A-positive ion mode and B-Negative ion mode) and LC/MS data of the identified compounds with their retention time, calculated m/z value and responses (frequency) were provided in **supplementary Table 1**. Identified compounds belonged to various classes including Flavonoids, Phenolic Acid, Alkaloids, Tri-terpenoids, Steroida, Fatty Acid, Amino Acids, Other Organic acids, Iridods and Vitamins. The major bioactive compounds were tabulated with their classification, therapeutic effect and references [26-104] (**Table 2**). The bioactive compounds have diverse therapeutic potential including anti– diabetic and anti–obesity activities.

Altogether, eleven different phytochemicals were identified by HPLC in the DMK extract namely, Gallic acid Vanillic Acid, Caffeic acid, Trigonelline, Adinosine, Rutin Quercetin Picriside I Pcroside II, Trans–Cinnamic acid and Apigenin (**Figure 2** and **Table 3**).

**Fig 2.**
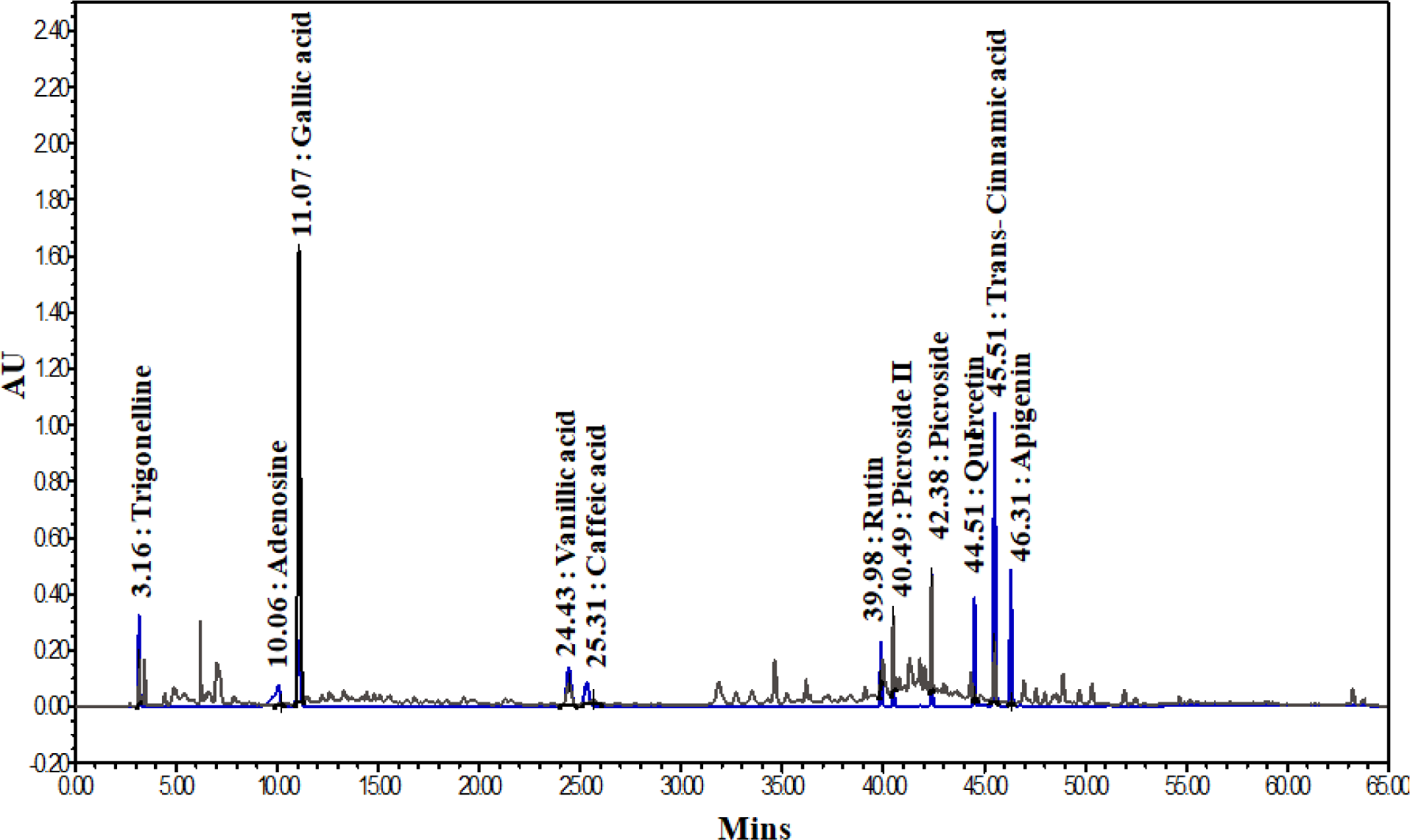
HPLC Profiling of the Major Constituents of Divya MadhuKalp (DMK). The HPLC chromatogram of the components was monitored at 230 nm. Here, the peaks showed for standard used in HPLC analysis are Gallic acid, Vanillic Acid, Caffeic Acid, Trigonelline, Adinosine, Rutin, Quercetin, Picroside I, Picroside II, Trans-cinnamic acid, Apigenin.

**Table 3.** Phenolics and marker compounds in Divya MadhuKalp (DMK) identified by HPLC analysis

| S.N. | Name of Marker Compound | Result (mg/gm) |
| --- | --- | --- |
| 1 | Gallic acid | 3.941 |
| 2 | Vanillic acid | 0.577 |
| 3 | Caffeic acid | 0.013 |
| 4 | Trigonelline | 0.319 |
| 5 | Adenosine | 0.063 |
| 6 | Rutin | 0.083 |
| 7 | Quercetin | 0.029 |
| 8 | Picroside I | 0.688 |
| 9 | Picroside II | 1.238 |
| 10 | Trans-Cinnamic acid | 0.162 |
| 11 | Apigenin | 0.004 |

### Effect of Divya MadhuKalp on Cell Viability

Treatment of 3T3-L1 cells with DMK and other extracts showed significant loss of cell viability at relatively high doses of 750 and 1000 µg/ml except *Trigonella foenum-graecum* (*Tf*). DMK and all the other constituent extracts were found to be non–toxic between the tested concentrations of 187.50 and 375 µg/ml in the 3T3-L1 cells. *Tf* treated 3T3-L1 cells showed ∼50% loss of cell viability at the high concentration of 93.75 µg/ml (**Figure 3**). L6 cells showed similar pattern of cell viability (data not shown).

**Fig 3.**
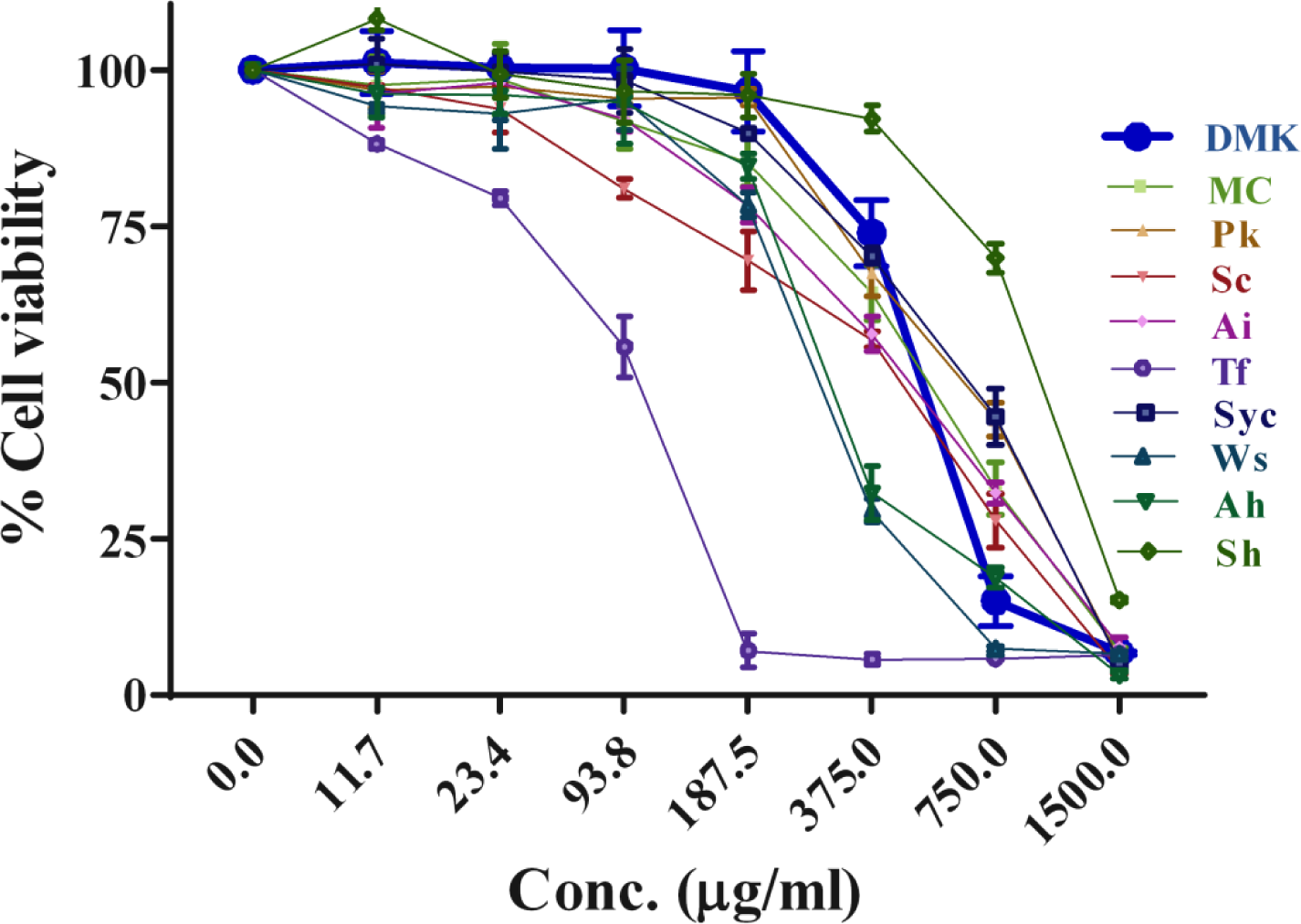
Effect of Divya MadhuKalp (DMK) and its Constituents on Cell Viability. The effects of DMK and its nine ingredients (*Mc*, *Pk*, *Sc*, *Ai*, *Tf*, *Syc*, *Ws*, *Ah*, and (*Mc*, *Pk*, *Sc*, *Ai*, *Tf*, *Sc*, *Wi, Ws*, *Ah*, and *Sh*) were studied for cell viability in mouse 3T3 L1 pre-adipocytes. Cell viability was determined by MTT after 24 h of treatment and extracts concentration ranged from 0-1500 µg/ml. Data represented the Mean ± SEM (n ≥3).

### Effect of Divya MadhuKalp on Glucose Uptake in L6 Cells

Efficacy of DMK and its nine components on cellular glucose uptake was studied in L6 cells using 2-NBDG, a fluorescent glucose analog. Insulin stimulated control L6 cells (Untreated) showed a 3 fold increase in the glucose uptake as compared to the insulin untreated (basal) cells. Co–treatment of the insulin treated L6 cells with DMK at the tested concentrations of 31.25, 62.5, 125 and, 250 μg/ml showed 3.6, 6.2, 7.3 and 7.8 fold increase in glucose uptake as compared to vehicle control under insulin stimulated condition (**Figure 4A**). Highest glucose uptake by the insulin stimulated L6 cells was measured at 250 μg/ml of DMK which was similar to reference drug Metformin (8.2 fold). Based on the dose response study, nine components of DMK were also screened independently at the concentration of 62.5 µg/ml for their glucose uptake potential. All the extracts showed significant increase in glucose uptake as compared to the vehicle control. Increase in glucose uptake were observed as 7, 3.8, 5, 5, 7.8, 7.5, 4.4, 7, 7.4, and 7.4 folds in DMK, *Mc*, *Pk*, *Sc*, *Ai*, *Tf*, *Syc*, *Ws*, *Ah* and Sh, respectively (**Figure 4B**).

**Fig 4.**
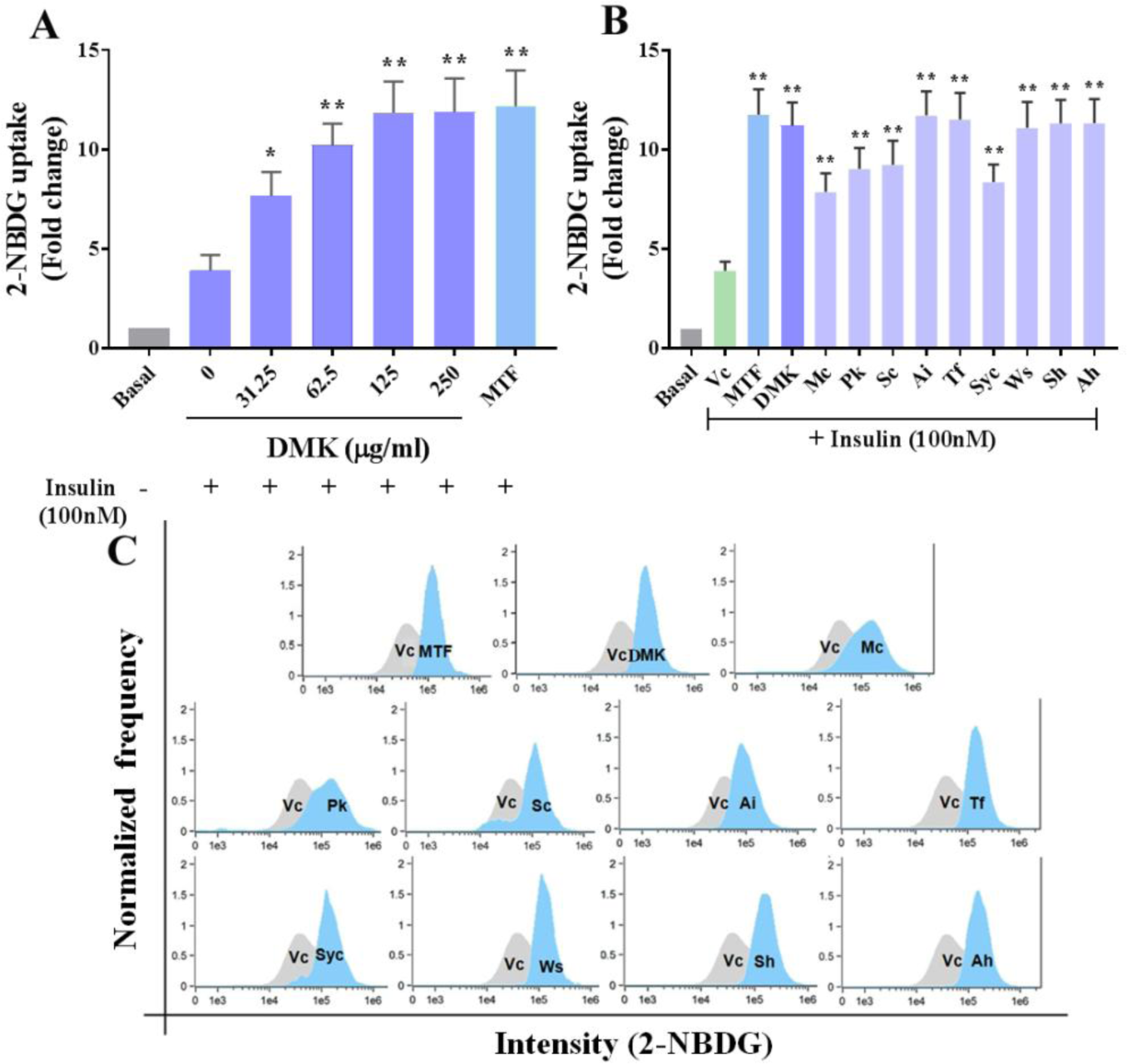
Effect of Divya MadhuKalp (DMK) and its Ingredients on Glucose Uptake in Rat Skeletal Muscle (L6) Cells. Cells were treated with extracts in presence of insulin for 24 h. Metformin (MTF– 100 µM) was used as a positive control. Glucose Uptake (2-NBDG 100 μg/ml) was quantified by Flow Cytometry of 8000 cells. (A) Effect of different doses of DMK (0-250 µg/ml) and (B) single dose (62.5 µg/ml) of its ingredients (*Mc*, *Pk*, *Sc*, *Ai*, *Tf*, *Syc*,*Sc*, *Wi, Ws*, *Ah*, and Sh) on glucose uptake. (C) Fluorescence intensity change (2-NBDG uptake) was observed in 3T3–L1 cells incubated with DMK and its ingredients (62.5 µg/ml). These data were harvested to compute the fold changes shown in (B). DMSO was used as a Vehicle control (Vc). All the data are expressed as Means ± SEM of three independent experiments. *p < 0.05 or **p< 0.01 as compared to vehicle control.

Flow cytometry analysis of treated cells showed changes in the fluorescence intensity of 2-NBDG, with respect to untreated cells. Increase in 2-NBDG intensity moved the peak maxima toward right in comparison to vehicle control and, this change in 2-NBDG intensity represents the glucose uptake potential of cells. **Figure 4C** showed the comparative glucose uptake of treated and untreated cells (Vc). These results demonstrated that DMK as well as its components are capable of stimulating glucose uptake under *in-vitro* conditions.

### Effect of Divya MadhuKalp on α-Glucosidase Activity and Advanced Glycation End Product (AGE) Formation

Inhibitors of α-glucosidase have been observed to delay glucose absorption by preventing carbohydrates digestion. In order to investigate the inhibitory effect of DMK extract, an *in-vitro* α-glucosidase inhibition test was performed. DMK displayed inhibitory activity for α-glucosidase enzyme with an IC_50_ value of 28.74 μg/ml. Acarbose was used as a positive reference control in a parallel experiment. DMK exhibited 55 % inhibition at 30 μg/ml and 97 % at 100 μg/ml concentration, respectively (**Figure 5A**).

**Fig 5.**
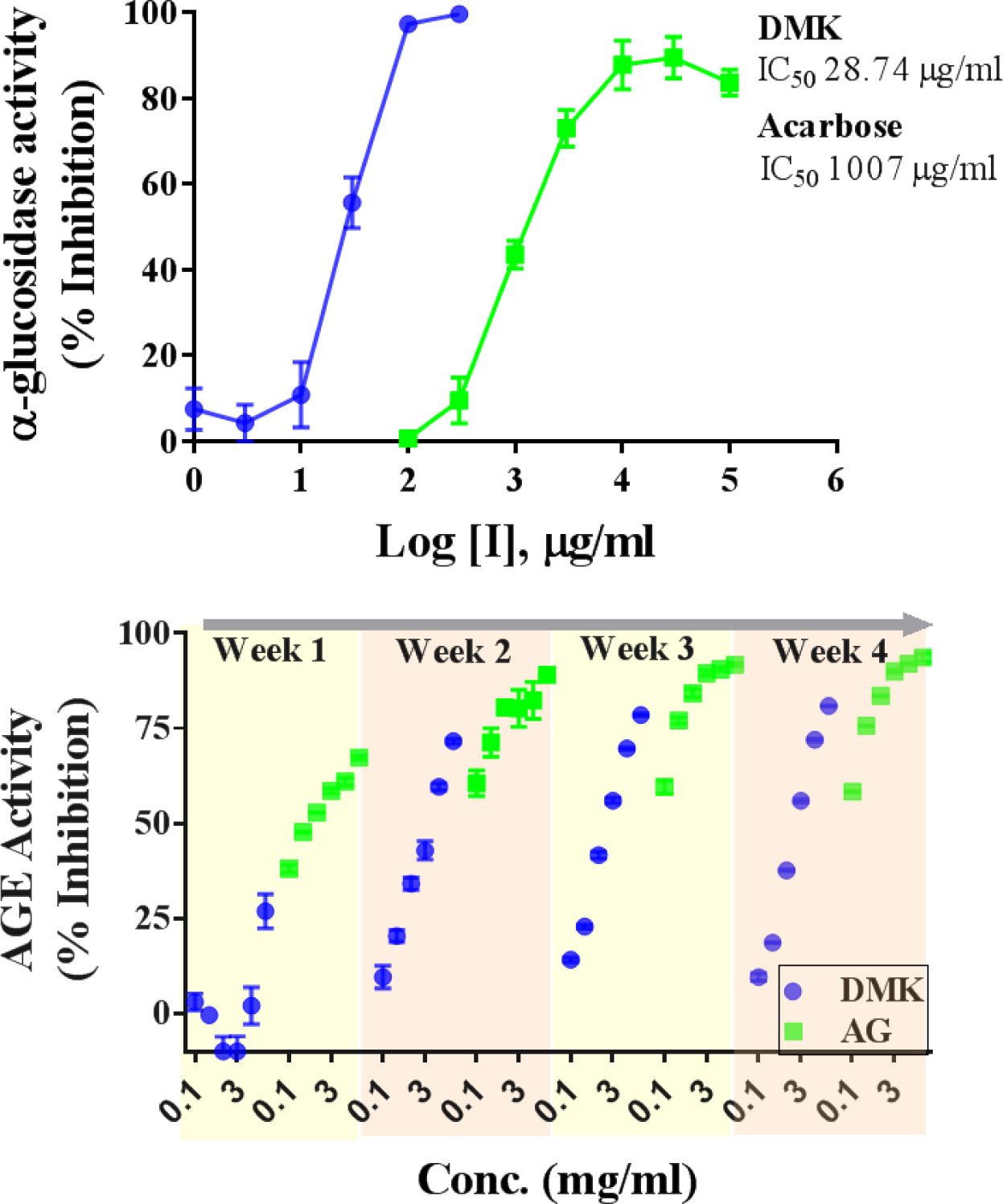
Effect of Divya MadhuKalp (DMK) on -Glucosidase and Anti Glycation Activity. (A) Inhibitory potency of DMK against-glucosidase activity. Acarbose is taken as the reference standard. (B) Effect on DMK Advanced Glycation End products (AGEs) of BSA up to 4 weeks of incubation. Aminoguanidine (AG) was used as the reference standard. The results were obtained from three independent experiments and expressed as Mean ± SEM.

The anti-glycation capacities of the DMK extracts was evaluated by the inhibition of the AGEs formation in the BSA/glucose system up to 4 weeks. The extract inhibited AGEs in a time and dose dependent manner. At 1^st^ week, the extract exhibited no significant inhibition where as from 2^nd^ −4^th^ week significant inhibitory capacity were observed at all concentrations compared to 1^st^ week. However, at 4th week inhibition was saturated with the extract of DMK. At 2^nd^ −4^th^ week a clear dose dependent (0.1 to 30 mg/ml) inhibition was observed. DMK exhibited a robust inhibitory response on AGEs formation with an IC50 value of 2.3 mg/ml, 1.4 mg/ml and 1.5 mg/ml whereas the positive control Aminoguanidine (AG) in the parallel experiments demonstrated the IC50 of 0.18 mg/ml, 0.08 mg/ml and 0.14 mg/ml in 2^nd^, 3^rd^ and 4^th^ week respectively (**Figure 5B**).

### Effect of Divya MadhuKalp on Blood Glucose Levels in Rats

DMK was tested for its anti–diabetic properties in animal model (Wistar rats) of hypoglycemia at the human equivalent dose of 200 mg/kg. Oral treatment with DMK (200 mg/kg) showed significant (p<0.01) reduction in blood glucose level (BGL) at 120 min as compared to the vehicle control treated animals (**Figure 6A**). Positive control, Glibenclamide at the tested concentration of 10 mg/kg exhibited significant decrease in BGL at 60 (p<0.01), 90 (p<0.01) and 120 min (p<0.01) as compared to the vehicle control animals.

**Fig 6.**
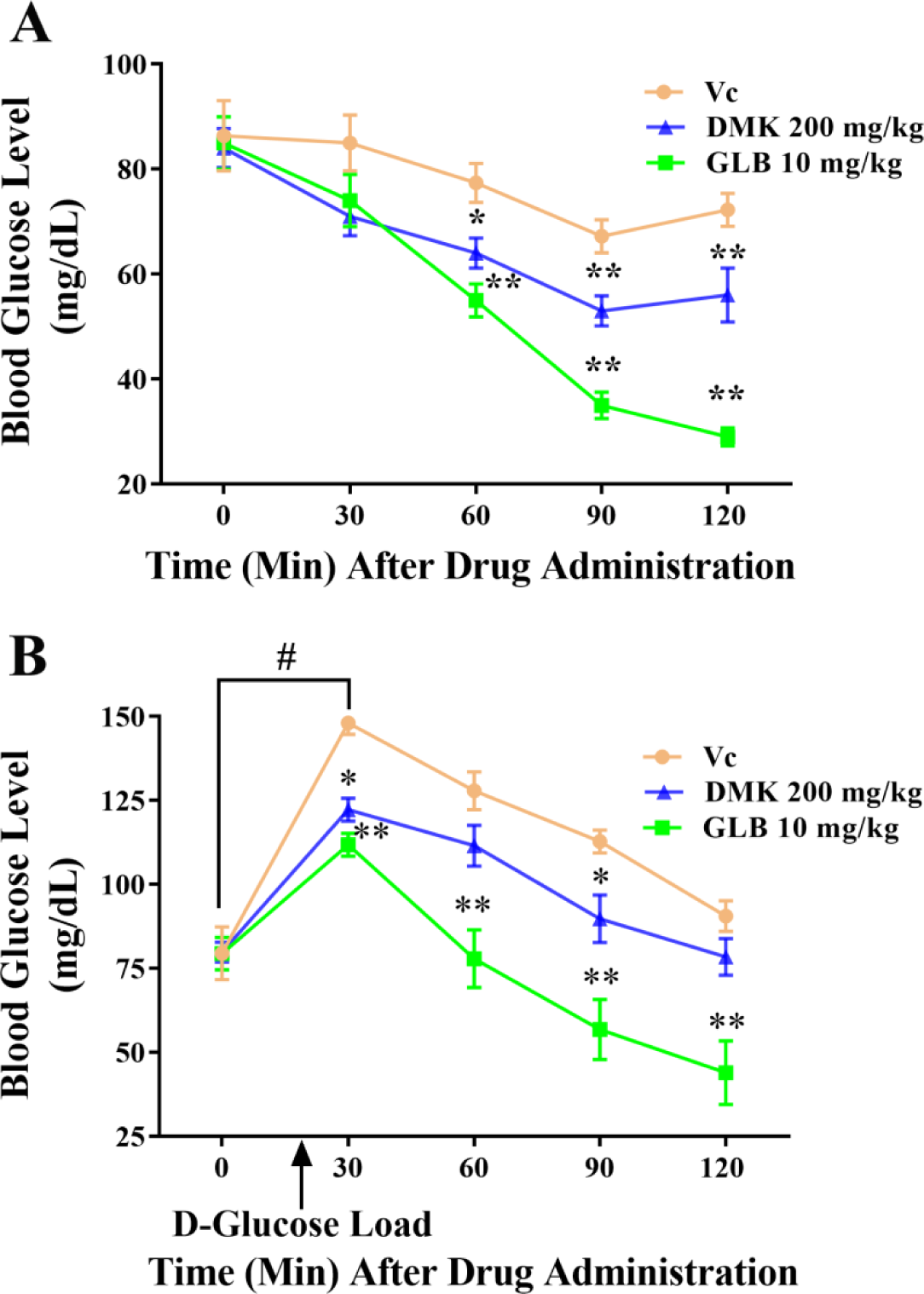
*In–Vivo* Efficacy of Divya MadhuKalp (DMK). (A) Effect of vehicle control (Vc), Glibenclamide (GLB; 10 mg/kg) and DMK (200 mg/kg) on blood glucose level in normal rats. (B) Effect of vehicle control (Vc), Glibenclamide (GLB; 10 mg/kg) and DMK (200 mg/kg) on blood glucose level of OGTT in normal rats. Values in the results are expressed as mean ± SEM, (n=8), # denotes significantly difference (p<0.05) in 0 min and 30 min time points (Student’s t–test); *p<0.05, **p<0.01 significantly different in comparison to control at respective time points (ANOVA followed by Dunnett’s t-test).

DMK was also tested for the Oral Glucose Tolerance Test (OGTT) in rats. Animals exhibited mean basal BGL of 79.6 ± 0.2 mg/dL after 12-14 h fasting. Vehicle–treated control animals dosed with D-glucose showed prominent (p<0.01) rise in BGL after 30 min, thereafter it returned to its baseline in 120 min (**Figure 6B**). Oral treatment of DMK at 200 mg/kg to glucose dosed animals showed significantly decreased BGL at 30 (p<0.05) and 90 min (p<0.05) as compared to control animals at these respective time points. In fact, the rate of BGL rise was also attenuated by DMK treatment, substantiating its glucose lowering properties in the intact animal systems. The standard of care drug, GLB at 10 mg/kg significantly lowered elevated BGL at 30 (p<0.01), 60 (p<0.01), 90 (p<0.05) and 120 min (p<0.01), as well (**Figure 6B**).

### Effects of Divya MadhuKalp on Adipocyte Differentiation

Anti-adipogenic activity of DMK and its nine constituents were evaluated in 3T3 L1 cells. Differentiation of pre-adipocytes into mature adipocytes was identified through the intracellular accumulation of lipid droplets in mature adipocytes, and quantified by Oil red O staining and, microscopy. In our study, 3T3 L1 cells treated with DMK extract gradually showed reduction of lipid droplets in a dose dependent manner (36.1% (31.2 μg/ml), 55.5 % (62.5 μg/ml), 67.9 % (125 μg/ml) and 80 % (250 μg/ml) (**Figure 7A and 7B**). Differentiated 3T3–L1 cells (vehicle control) showed enhanced lipid accumulation (∼65 %) when compared with the pre-adipocytes (UI). Investigation of the anti–adipogenic potential of individual components of DMK revealed the *Tf* (67.4 %), *Ws* (65.4 %) and *Syc* (50.55 %) to exhibit significant inhibition in lipid accumulation as compared to the vehicle control. Microscopic observations also revealed that cells treated with DMK, *Tf*, *Ws* and *Syc* maintained fibroblastic shape and contained less lipid droplets (**Figure 7C and 7D**).

**Fig 7.**
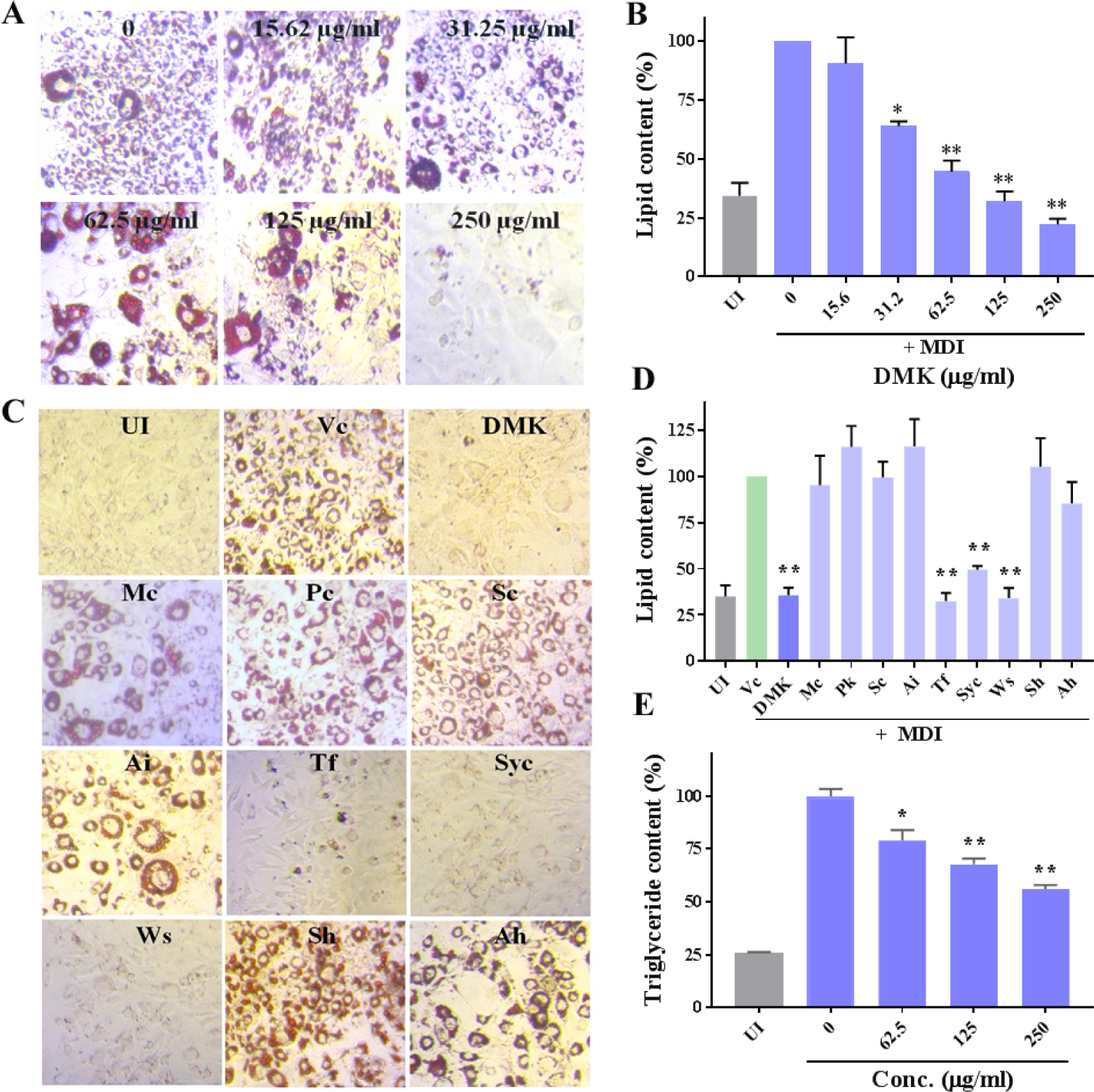
Anti-adipogenic Effect of Divya MadhuKalp (DMK) and its Ingredients in Mouse 3T3 L1 Pre-adipocytes. Lipids were stained with Oil Red O (MDI: adipogenic differentiation media) and quantified at 492 nm. (A) Representative photomicrographs at 20X showed lipid accumulation in the cells and, (B) Relative lipid content of DMK (0-250 µg/ml) treated cells. (C) Microscopic images (20X) of Oil red O stained adipocytes treated with DMK ingredients (*Mc*, *Pk*, *Sc*, *Ai*, *Tf*, *Syc*,*Sc*, *Ws*, *Ah*, and *Sh*) (D) Relative lipid content of DMK ingredients. The dose used for each ingredient was 250 µg/ml except for *Tf* (62.5 µg/ml) based on MTT assay. (E) Effect of DMK on the triglyceride deposition in differentiated 3T3–L1 cells. Cells were cultured in adipogenic differentiation media with or without treatment of DMK and, triglyceride content was quantified at 540 nm. DMSO was used as a Vehicle control (Vc). Uninduced (UI) cells were not treated with MDI. All the data were expressed as means ± SEM of three independent experiments. UI: Uninduced. *p < 0.05 or **p< 0.01 as compared to Vehicle control (Vc).

Intracellular quantification of the triglycerides content on day 10 of adipogenic differentiation showed significant reduction in triglycerides accumulation following DMK treatment in a dose–dependent manner [62.5 µg/ml (21.1 %), 125 µg/ml (32.26 %) and 250 µg/ml (43.85 %)] (**Figure 7E**). Taken together, these finding strongly suggest that DMK inhibits adipogenesis differentiation in 3T3 L1 pre–adipocytes.

### Effect of Divya MadhuKalp on Expression of Adipogenic Transcription Factors and Marker Genes

Influence of DMK on the mRNA expressions of transcription factors; PPARγ and C/EBPα involved in adipocyte differentiation was studied in 3T3 L1 cells. In our study, the expression level of PPARγ was inhibited at the DMK concentrations of 125 μg/ml (3.2 fold) and 250 μg/ml (4.1 fold), respectively. Similarly, C/EBPα showed 3.53 and 4.7 fold inhibition in expression level at the same tested DMK concentrations (**Figure 8A & 8B**).

**Fig 8.**
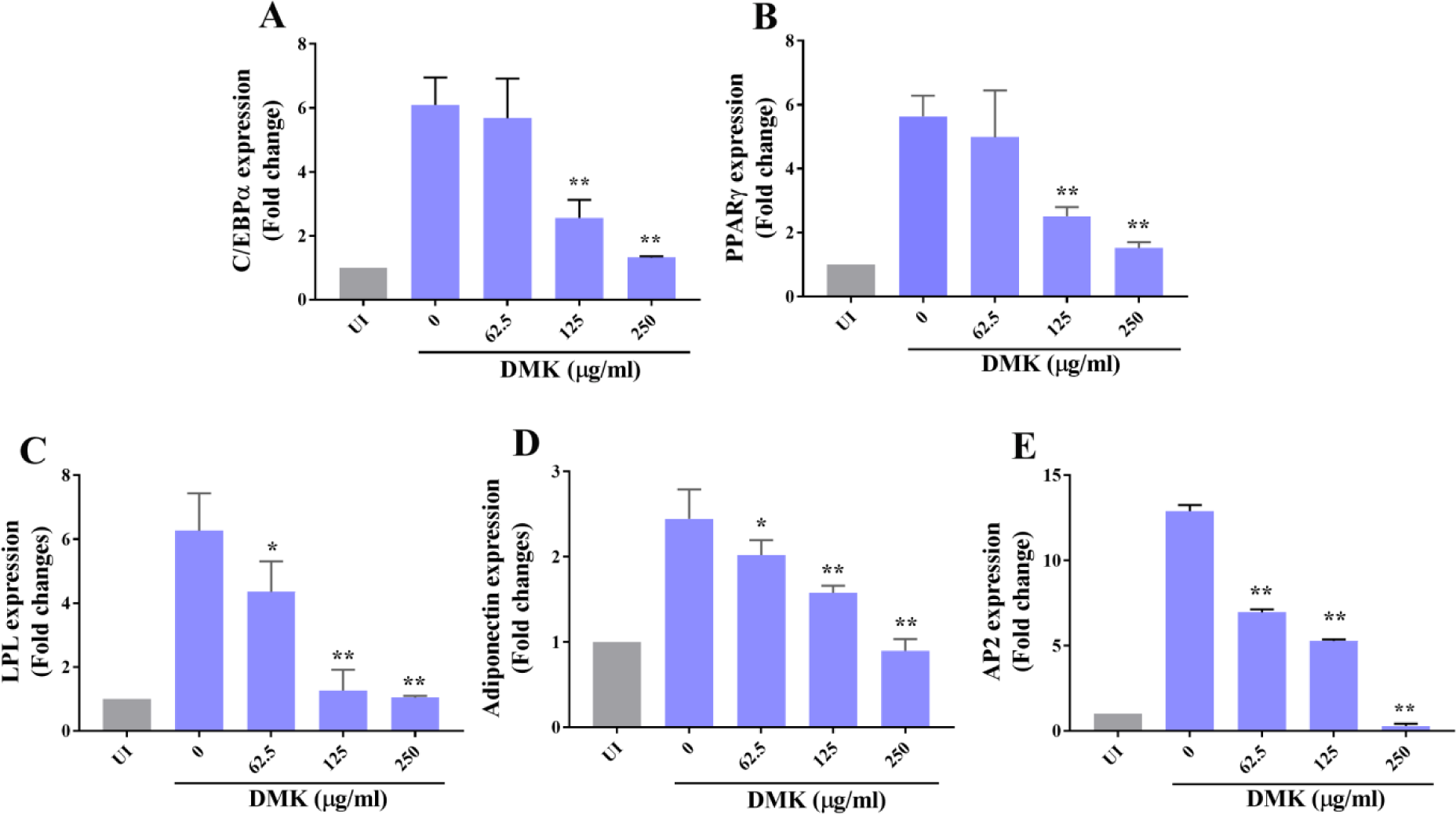
Effect of Divya MadhuKalp (DMK) on the Expression of AdipogenicTranscription Factor and Target Gene Markers. The 3T3 L1 mouse pre-adipocytes were induced to differentiate into adipocytes in MDI (adipogenic differentiation media) medium with (62.5, 125 and 250 µg/ml) or without DMK. Uninduced (UI) cells were not treated with MDI. mRNA was extracted and the expression of adipogenic transcription factors and related adipogenic modulators genes were detected using one step RT-PCR. The relative expression level of (A) PPARγ, (B) C/EBPα, (C) LPL, (D) AP2, and (E) Adiponectin are presented as the fold change in DMK treated groups vs. untreated after normalization to GAPDH mRNA levels. Values are expressed as the mean ± SEM. *p < 0.05 or **p< 0.01 as compared to control (Untreated).

Further downstream gene expression profiling of the adipocyte specific genes such as adiponectin, LPL and AP2 was done in DMK treated 3T3 L1 cells. Downregulation of LPL gene was noted with 2, 5 and 5.23 fold at 62.5, 125 and 250 µg/ml concentrations, respectively (**Figure 8C**). Similarly, expression of AP2 was inhibited significantly by all tested concentrations of DMK, showing a 6, 7.6 and 12.8 fold inhibition effect at 62.5, 125 and 250 µg/ml concentrations, respectively (**Figure 8D**). The fold inhibition in the expression level of adiponectin was found at 125 μg/ml (1.2 fold) and 250 μg/ml (2 fold) concentrations of DMK compared to untreated cells (**Figure 8E**). These results suggested that DMK regulated the lipid accumulation by downregulating expression of adipogenic genes, resulting in the prevention of adipogenesis.

## Discussion

Plants remain as an important source of therapeutic material since antiquity. Several studies have been conducted to find a safe and effective therapy to treat diabetes in traditional medicine system^105,106,107^. Standardization of polyherbal formulation is required to evaluate the quality and safety of herbal product to use it as an authentic drug^19, 108^. In the present study, DMK extract was evaluated to identify the presence of bioactive chemical compounds using LC/MS analysis. The result showed that total 139 compounds including flavonoids, phenolic acids, terpinoids, phytosteroids, fatty acids, iridoid glycosides, vitamins and amino acids are present in the extract. The analysis also revealed that flavonoids and organic acids are the major compounds occurring in DMK extract. Therefore, major therapeutic activities might come from the flavonoids and organic acids present in the DMK.

Diabetes is characterized by high blood sugar levels, therefore lowering blood sugar levels in hyperglycemic state is the main treatment method in diabetes and its associated complications. Skeletal muscle is the major target site for insulin stimulated glucose uptake and plays an important role in postprandial glucose regulation^109^. In the present study, we observed that DMK enhanced glucose uptake significantly in skeletal muscle (L6) cells, even the lowest tested concentration (31.25 µg/ml) showed 3.6 fold increase in cellular glucose uptake as compared to vehicle control in insulin–stimulated condition. Hypoglycemic activity of DMK was also evaluated in normoglycemic animals. The results demonstrated that a significant reduction in blood glucose level at human equivalent (200 mg/kg) dose of DMK. In addition, OGTT was used to detect the impaired glucose tolerance that represents pre-diabetic and diabetic conditions. Our results displayed that the treatment of DMK at human equivalent dose prominently modulated glucose tolerance in normal rats by minimizing the blood glucose level (BGL) peak post glucose loading. These studies indicated that the increased glucose uptake or hypoglycemic effect in the target cells or tissue due to the synergistic effect of various bioactive compounds present in DMK which probably stimulated the pancreatic β-cells to produce insulin and its sensitization or possesses an insulin like or extra-pancreatic mechanism of action (Table 3).

One of the therapeutic approaches to regulate postprandial glucose level is inhibition of carbohydrate hydrolyzing enzymes, such as α-glucosidase^41^. DMK has shown considerable α-glucosidase inhibition under cell free conditions and this inhibition might be due to the synergistic action of phenols, flavonoids and other compounds present in DMK. In addition, DMK demonstrated a significant anti-glycation activity under cell free condition. AGEs are proteins or lipids that become glycated because of long standing sugar exposures in diabetes and, contribute in the progression of diabetes associated complications. The anti-glycation activity was found to be correlated with flavonoids, phenolic acids and other phytochemicals of the DMK. The enhanced cellular glucose uptake and anti-alpha glucosidase activity of DMK provides the evidence in support of its anti-diabetic potential. Moreover, the anti-glycation activity indicates the potential of DMK in treating diabetic associated complications.

Obesity is closely associated with diabetes mellitus and characterized by excessive growth of adipose tissue mass because of hyperplasia and hypertrophy of adipocytes. Inhibition of pre-adipocytes differentiation is one of the key therapeutic strategies to inhibit adipogenesis^3,4,5^. Our results showed that DMK significantly suppressed the adipogenic differentiation, lipid content and intracellular triglycerides in 3T3 L1 cells. A significant reduction in lipid content (80 %) and triglyceride content (43.85 %) was noted at 250 µg/ml concentration of DMK. Adipogenesis involves the process of differentiation of pre–adipocytes into adipocytes and it tightly synchronized with the sequential activations of various transcriptional regulators and downstream genes^4, 5^. A downregulation observed at mRNA expression of two important transcription master regulators (PPARγ and C/EBPα) of adipogenesis indicated anti–adipogenic potential of DMK. Furthermore, we demonstrated that DMK downregulated mRNA expression of downstream adipogenic marker (LPL, AP2, and adiponectin) genes that are responsible for lipid accumulation in adipocytes. LPL plays a critical role in triglyceride uptake and storage. Adiponectin plays an important role in enhancing insulin sensitivity and increasing fatty acid oxidation in liver and muscle. In addition to adiponectin, AP2 is also an adipokines have a specific role in lipolysis^110^. Therefore, the bioactive phytochemicals in DMK might have capacity to inhibit adipocyte differentiation by down regulating transcription factors and adipogenic genes that could be an effective strategy in the treatment of obesity.

Several studies reported that medicinal property of various plants was correlated with their bioactive phytochemical compounds which have therapeutic potential against diseases ^7, 105, 106, 107^. The herbo-mineral formulation DMK contains multiple bioactive compounds. The understanding of mode of action of these bioactive compounds on multiple targets is very important for cure and prevention of a disease. These compounds have been studied extensively with their chemical structure and mode of action. Catechin, a flavonoid present in DMK exhibit anti-hyperglycemic, anti-hyperlipidemic, anti-glycosidase and anti-AGEs activities by enhancing anti-oxidant defense system ^26, 27, 28^. Myricetin demonstrated glucoregulatory activity as a GLP-1R agonist for the treatment of T2DM ^29^. Also it suppresses differentiation of pre-adipocytes into adipocytes by down regulating transcription factor (PPARγ, C/EBPα) and adipogenic marker genes, in addition, it enhances lipolysis by releasing glycerol from fully differentiated adipocytes^30^. DMK contains Astragalin which is multifaceted compound regulating various molecular targets such as transcription factors, enzymes, kinases etc. with diversified pharmacological applications including anti-obesity and anti–diabetic properties ^31^. The flavonoid rutin has anti– hyperglycemic property including decrease of carbohydrates absorption in small intestine, increase of tissue glucose uptake, stimulation of insulin secretion from beta cells, protecting Langerhans islet against degeneration and inhibition of tissue gluconeogenesis ^32^. Rutin also decreases reactive oxygen species, advanced glycation end-product precursors and prevent all kind of pathologies associated with diabetes ^33^. DMK contain Quercetin, which is a flavonoid improving insulin secretion and sensitizing activities as well as glucose utilization in peripheral tissues ^35^. It inhibits the formation of AGEs by blocking reactive dicarbonyl compounds, identified as its precursors ^36^. Also, it inhibits the adipogenesis of muscle satellite cells *in-vitro* by suppressing the transcription of adipogenic markers ^37^. Jung et al demonstrated that apigenin significantly reduced levels of fasting blood glucose, hyper-insulinemia and HOMA-IR, a surrogate marker for insulin resistance, in type I and type II diabetic animal. Moreover, it significantly decreased hepatic PEPCK and G6Pase activity controlling hepatic gluconeogenesis which is closely related to diabetes and obesity ^39^. Vitexin and isovitexin are naturally occurring C-glycosylated derivatives of apigenin. Both compound showed highly anti α-glucosidase inhibitory activity ^41^. Also, two compounds inhibit the formation of AGEs significantly but failed to trap reactive carbonyl species, the anti AGEs activities may be due to their free radical scavenging capacity ^42^. Vitexin prevented HFD induced obesity/adipogenesis via the AMPKα mediated pathway ^43^. Kaempferol decreased TG accumulation at the dose of 25 µM by reducing the transcription factor C/EBPα in human mesenchymal stem cells (MSCs) ^39^. Hyperoside significantly inhibited receptor for anti–glycation end product (RAGE) expression and promoted proliferation in AGE–stimulated ECV304 cells ^45^. Daidzein stimulated glucose uptake in L6 by AMPK phosphorylation increasing GLUT4 translocation to PM of muscle cells and significantly suppressed the rises in blood glucose levels in db/db and KK–Ay mice ^46^. Also it induced survival of pancreatic β–cells and insulin secretion in type1 diabetic animal model ^47^. Luteolin exerted anti-adipogenic effects by down regulating adipogenic transcription factors, suppressed of NF-κB and MAPKs activation, and also improved insulin sensitivity in 3T3 L1 adipocytes, suggesting that luteolin prevents obesity, associated inflammation and insulin resistance ^49, 50^. Vicenin 2 significantly exhibited anti–diabetic activity by inhibition of α-glucosidase and protein tyrosine phosphatase 1B (PTP1B). It also inhibited AGEs formation by attenuating the formation of protein carbonyl groups as well as modification of protein thiol groups ^56^. Baicalein play a novel anti-diabetic action by directly improving β-cell and human islet viability and insulin secretion ^57^. It decreased the intracellular lipid accumulation by down–regulation of glucose uptake via repression of Akt-C/EBPα-GLUT4 signaling ^58^. Chlorogenic acid reduces islet cell apoptosis and improves pancreatic function ^64^, also it improves glucose and lipid metabolism, via AMPK activation ^65^. It lowered the levels of fasting plasma glucose and HbA1c during late diabetes in db/db mice ^66^. Apocynin has the potential to improve insulin sensitivity ^68^. Ferulic acid improved insulin sensitivity and hepatic glycogenesis but inhibits gluconeogenesis to maintain normal glucose homeostasis in type 2 diabetic rat ^69^. Also, it has a potential to modulate adipogenesis via expressing of heme oxygenase-1 (HO-1) and suppressing C/EBPα and PPARγ, triglyceride–synthesizing enzymes, fatty acid synthase (FASN) and acetyl-CoA carboxylase (ACC) ^70^. Caffeic acid esters are able to stimulate glucose transport in skeletal muscle via increasing GLUT4 translocation, reduce hepatocellular glucose production and adipogenesis in an AMPK-dependent manner ^71^. Ellagic acid lowered significantly blood glucose level by stimulating insulin secretion ^73^. Gallic acid and its derivatives play significant role through activation of the AMPK/Sirt/PGC1α pathway ^74^. Furthermore, it exhibited a strong protective effect on disease progression in streptozotocin (STZ) injected type 1 and 2 diabetic models. Therefore, Gallic acid has a potential therapeutic intervention in the protection and/or improvement of metabolic syndromes ^75, 76, 78, 79^. Stearidonic acid inhibited adipocyte differentiation and reduce lipid accumulation in 3T3 L1 adipocytes through down–regulation of adipogenic transcription factors and adipogenic genes ^94^. Withaferin A is a leptin sensitizer; its treatment showed 20–25% reduction of body weight of diet–induced obese mice and also decreased obesity–associated abnormalities including hepatic steatosis. Withaferin A also affects the body weight of *ob/ob* and *db/db* mice which are deficient in leptin signaling suggesting that it has leptin independent effects on glucose homeostasis ^97^. Diosgenin a steroid saponin showed hypo-glycaemic effect and improved dyslipidemia by decreasing the hepatic lipid content in type-II diabetic model ^103^. The synergistic effect of phyto-constituents present in polyherbal formulation always acts on multiple target and pathways of disease, which is not available in single herb formulation ^8, 132^. Moreover, polyherbal formulation contains very minimum amount of compound exert very little or no side effect compared to single herb formulation. The enhanced effectiveness of the DMK as anti diabetic and anti adipogenic agent might be through additive or synergistic actions of its multiple bioactive compounds.

We concluded that the DMK is capable of enhancing glucose uptake in cells, lowering blood sugar level, anti-α glucosidase activity, anti-glycation activity and anti-adipogenic property indicating its potential in treating diabetic and its associated complication. All the activities of DMK could be due to the synergistic effect of bioactive compounds. Moreover, no side effect is expected as the formulation is fully nature derived and has long term historical use. The observed multi targets mode of action of DMK puts it in a rather different class of therapeutic agents; and offers a new preventive and therapeutic option in holistic DM management.

## Acknowledgments

We are thankful to Patanjali Research Foundation Trust, Haridwar, India for financial support. We thank Mr. Sunil Kumar and Pratima Kumari for their valuable help with culturing cells and extracts preparation. We are also thankful to Mr. Lalit Mohan, Mr. Tarun Ji, and Mr.Gagan Ji for their administrative support.

## Author Contributions

Conceptualization and experimental design: SKD, AJ, VS, CSJ, AB. Performed the experiments: SKD, AJ, LB, KJ, VKS, NS, SV Analyzed the data: SKD, AJ, SS, VS, CSJ, Manuscript writing: AJ, SKD, VS, SS, Manuscript reviewed: SKD, AJ, VS, CSJ, VKS. All authors read and approved the final version of the manuscript.

