## Supplementary Table for "LC/MS-QToF Profiling, Anti-Diabetic and Anti-Adipogenic potential of Divya MadhuKalp: A Novel Herbo-mineral Formulation"

**Supplementary Table 1. LC/MS-QToF analysis for the phytochemicals screening of hydroalcoholic extract of Divya MadhuKalp (DMK)**

| **S.N.** | **Component  name** | **Positive ion mode** | | | **Negative ion mode** | | |
| --- | --- | --- | --- | --- | --- | --- | --- |
|  |  | **Observed RT (min)** | **Observed m/z** | **Response** | **Observed RT (min)** | **Observed m/z** | **Response** |
| **1** | Arginine | 0.54 | 175.119 | 45435 | 0.55 | 173.1042 | 8581 |
| **2** | Aspartic acid |  |  |  | 0.58 | 132.0306 | 9583 |
| **3** | Galactomannan |  |  |  | 0.58 | 503.1616 | 47137 |
| **4** | Glutamic acid | 0.6 | 148.0602 | 5617 |  |  |  |
| **5** | Sucrose |  |  |  | 0.6 | 341.1079 | 66366 |
| **6** | Valine | 0.62 | 118.0865 | 441415 |  |  |  |
| **7** | Trigonelline | 0.63 | 138.0551 | 365080 |  |  |  |
| **8** | Cytosine | 0.64 | 112.05 | 9707 |  |  |  |
| **9** | Leucine | 0.65 | 132.1017 | 8340 |  |  |  |
| **10** | Pseudotropine | 0.66 | 142.1225 | 34885 |  |  |  |
| **11** | Malic acid |  |  |  | 0.7 | 133.014 | 248831 |
| **12** | Pellentierine | 0.85 | 142.1224 | 18917 |  |  |  |
| **13** | Nicotinic acid | 0.9 | 124.0394 | 25233 |  |  |  |
| **14** | Citric acid | 0.92 | 191.0197 | 723676 |  |  |  |
| **15** | Uracil | 0.99 | 113.0348 | 14135 |  |  |  |
| **16** | Tyrosine | 1.07 | 182.0813 | 55119 | 1.08 | 180.0662 | 13877 |
| **17** | Adenosine | 1.13 | 268.105 | 154269 |  |  |  |
| **18** | Ascorbic acid | 1.17 | 175.0243 | 29835 |  |  |  |
| **19** | Isoleucine | 1.18 | 132.1017 | 10215 |  |  |  |
| **20** | Guanosine | 1.27 | 284.1006 | 72046 | 1.28 | 282.0843 | 54153 |
| **21** | Gallic acid | 1.62 | 171.0285 | 146444 | 1.63 | 169.0139 | 1932468 |
| **22** | Phenylalanine | 1.92 | 166.0858 | 97766 | 1.93 | 164.071 | 19115 |
| **23** | Fulvic acid | 2.81 | 309.0599 | 14813 |  |  |  |
| **24** | Protocatechuic acid | 2.96 | 153.0189 | 82789 |  |  |  |
| **25** | Ethyl gallate | 2.97 | 197.045 | 15418 |  |  |  |
| **26** | Hetisinone | 3.27 | 328.1903 | 17807 |  |  |  |
| **27** | Tryptophan | 3.63 | 205.0957 | 14748 | 3.64 | 203.0816 | 7620 |
| **28** | Methyl gallate | 5.03 | 183.0296 | 66564 |  |  |  |
| **29** | 2,6-Di-O-galloyl-β-D-glucose | 5.26 | 485.0927 | 28454 | 5.27 | 483.0783 | 496218 |
| **30** | Heteratisine | 5.42 | 392.243 | 37170 |  |  |  |
| **31** | Swertiamarin |  |  |  | 5.51 | 373.1145 | 310043 |
| **32** | 3,6-Di-O-Galloyl-β-D-glucose | 5.75 | 483.0785 | 572851 |  |  |  |
| **33** | Catechin |  |  |  | 5.84 | 289.0708 | 5358 |
| **34** | Chlorogenic acid | 5.86 | 355.1026 | 151252 | 5.88 | 353.0883 | 509024 |
| **35** | Hetisine | 5.94 | 330.2067 | 302443 |  |  |  |
| **36** | Naringenin-7-O-glucoside | 6.02 | 435.1285 | 64689 | 6.04 | 433.1146 | 145827 |
| **37** | Daidzin | 6.02 | 417.1179 | 8041 |  |  |  |
| **38** | Vanillic acid | 6.04 | 169.0493 | 71019 | 6.05 | 167.0351 | 48905 |
| **39** | Atidine | 6.34 | 360.2522 | 10233 |  |  |  |
| **40** | Caffeic acid |  |  |  | 6.47 | 179.0347 | 54434 |
| **41** | Enicoflavine |  |  |  | 6.7 | 210.0766 | 28395 |
| **42** | Dihydrophaseic  acid |  |  |  | 7.61 | 281.1397 | 36872 |
| **43** | Corilagin |  |  |  | 7.86 | 633.0744 | 2493819 |
| **44** | Irilone | 7.88 | 299.0551 | 9978 |  |  |  |
| **45** | Mangiferin | 7.88 | 423.0926 | 684156 | 7.9 | 421.0785 | 879898 |
| **46** | Riboflavin | 8.03 | 377.1454 | 45794 | 8.04 | 375.1304 | 6954 |
| **47** | Gentiopicrin | 8.27 | 357.1182 | 7269 |  |  |  |
| **48** | Vicenin 2 | 8.34 | 595.167 | 150513 | 8.34 | 593.1517 | 169657 |
| **49** | Atisine | 8.49 | 344.2588 | 571091 |  |  |  |
| **50** | Sweroside | 8.59 | 357.1178 | 6589 |  |  |  |
| **51** | Vicenin 1 | 9.05 | 565.1558 | 643291 | 9.06 | 563.1407 | 806694 |
| **52** | Apocynin | 9.13 | 167.0703 | 83240 |  |  |  |
| **53** | Vicenin 3 | 9.35 | 565.1562 | 604235 | 9.36 | 563.1407 | 782796 |
| **54** | Myricetin | 9.35 | 319.0453 | 39417 |  |  |  |
| **55** | Quercetin-3-galactoside-7-glucoside | 9.35 | 627.157 | 97432 | 9.36 | 625.1408 | 348797 |
| **56** | Astragalin | 9.39 | 449.1084 | 325471 | 9.4 | 447.0937 | 395613 |
| **57** | Quercetin-6-O-glucoside | 9.52 | 481.0987 | 36477 | 9.53 | 479.0838 | 139007 |
| **58** | Scopoletin | 9.53 | 193.0492 | 12722 |  |  |  |
| **59** | Kaempferol-3-glucuronide | 9.69 | 465.1025 | 5443 | 9.69 | 463.0869 | 9425 |
| **60** | Ferulic acid | 9.69 | 195.0648 | 13901 | 9.71 | 193.0501 | 33779 |
| **61** | Kutkin | 9.71 | 497.1637 | 61617 |  |  |  |
| **62** | Vitexin | 10.34 | 433.1133 | 487845 | 10.34 | 431.0987 | 575635 |
| **63** | Calycosin |  |  |  | 10.34 | 283.0603 | 10612 |
| **64** | Rutin | 10.49 | 611.1618 | 723121 | 10.49 | 609.1458 | 1565541 |
| **65** | Quercetin-3-O-glucoside | 10.49 | 465.1031 | 64868 | 10.48 | 463.0881 | 68070 |
| **66** | Isovitexin | 10.54 | 433.1135 | 658305 | 10.55 | 431.0986 | 377274 |
| **67** | Ellagic acid |  |  |  | 10.56 | 300.9991 | 1250890 |
| **68** | Picroside II | 10.64 | 513.1609 | 75365 | 10.65 | 511.1456 | 2080186 |
| **69** | Gentiopicrin |  |  |  | 10.68 | 355.1035 | 100597 |
| **70** | Hyperoside | 10.81 | 465.1032 | 210602 | 10.82 | 463.0881 | 791174 |
| **71** | ViscumneosideⅡ | 10.82 | 535.1455 | 174070 | 10.82 | 533.13 | 222938 |
| **72** | Kaempferol 7-O-glucoside | 10.9 | 449.1068 | 22041 | 10.91 | 447.0924 | 27612 |
| **73** | Daidzin |  |  |  | 11.07 | 415.1029 | 14983 |
| **74** | β-Hydroxyacteoside |  |  |  | 11.25 | 639.1925 | 1663872 |
| **75** | 3,5-Dicaffeoylquinic acid | 11.49 | 517.134 | 52001 | 11.5 | 515.1195 | 379673 |
| **76** | Kaempferol-3-O-rutinoside | 11.53 | 595.166 | 146007 | 11.54 | 593.1504 | 414397 |
| **77** | Azadirachtol |  |  |  | 11.83 | 579.2088 | 113609 |
| **78** | Tricin | 12.14 | 331.0812 | 11303 |  |  |  |
| **79** | Baicalin |  |  |  | 12.27 | 445.0779 | 506718 |
| **80** | Picroside I | 12.54 | 493.1708 | 49464 |  |  |  |
| **81** | Picroside III |  |  |  | 12.55 | 537.1636 | 2765621 |
| **82** | Minecoside | 12.95 | 539.176 | 34595 | 12.96 | 537.1618 | 1363034 |
| **83** | Kaempferol-3-O-β-D-glucuronide | 13.55 | 463.0872 | 109392 | 13.56 | 461.0744 | 122056 |
| **84** | Luteolin | 13.55 | 287.0547 | 32377 | 13.56 | 285.0401 | 29765 |
| **85** | Kutkoside |  |  |  | 13.94 | 511.1464 | 158965 |
| **86** | Veronicoside |  |  |  | 13.95 | 465.1393 | 12568 |
| **87** | Orientin-2-o-p-trans-coumarate | 13.97 | 595.1454 | 183940 | 13.98 | 593.1301 | 177099 |
| **88** | 1,3,7-Trimethoxyxanthone | 14.2 | 287.0914 | 283853 | 14.21 | 285.0773 | 186718 |
| **89** | Blumenol A |  |  |  | 14.28 | 223.1334 | 6965 |
| **90** | Cinnamic acid | 14.34 | 149.0591 | 21117 |  |  |  |
| **91** | Quercetin | 14.93 | 303.0503 | 209548 | 14.93 | 301.0355 | 352949 |
| **92** | Kaempferol | 14.96 | 287.0557 | 46685 | 14.97 | 285.0404 | 53627 |
| **93** | Rhaponticin |  |  |  | 15.08 | 419.1348 | 68860 |
| **94** | Withanoside IV | 16 | 783.4173 | 10935 |  |  |  |
| **95** | Trigofoenoside A | 16.69 | 903.4959 | 24868 |  |  |  |
| **96** | Gitogenin | 16.79 | 433.3304 | 12986 |  |  |  |
| **97** | Apigenin | 16.83 | 271.0595 | 49045 | 16.84 | 269.0458 | 78550 |
| **98** | 5,7-Dihydroxy-4'-methoxyisoflavanone | 17.49 | 287.0911 | 161021 | 17.5 | 285.077 | 494756 |
| **99** | Yamogenin | 17.56 | 415.3201 | 10153 |  |  |  |
| **100** | Tigogenin-3-O-β-D-glucopyranosyl(1→4)-β-D-galactopyranoside | 17.58 | 741.4433 | 194892 |  |  |  |
| **101** | Smilagenin | 17.68 | 417.3353 | 19843 |  |  |  |
| **102** | 1,3,7,8-Tetrahydroxyxanthone | 18.2 | 261.0393 | 61847 | 18.21 | 259.0248 | 525906 |
| **103** | Momordicoside L | 18.28 | 635.4159 | 115753 |  |  |  |
| **104** | Naringenin |  |  |  | 18.3 | 271.0613 | 124167 |
| **105** | Sugiol | 18.44 | 301.2162 | 116318 |  |  |  |
| **106** | Trillin | 18.65 | 577.3737 | 185264 |  |  |  |
| **107** | Diosgenin | 18.65 | 415.3199 | 31128 |  |  |  |
| **108** | Withanolide E |  |  |  | 19.45 | 485.2522 | 11728 |
| **109** | Withaferin A | 19.47 | 471.2747 | 36566 |  |  |  |
| **110** | Withanoside V | 20.2 | 767.4208 | 9747 |  |  |  |
| **111** | Withanolide D | 20.54 | 471.2742 | 34991 |  |  |  |
| **112** | Withanolide A | 21.34 | 471.2738 | 41444 |  |  |  |
| **113** | Swertianin | 22.29 | 275.0551 | 194206 | 22.3 | 273.0404 | 451927 |
| **114** | Tigogenin | 22.92 | 417.3353 | 14488 |  |  |  |
| **115** | 1-Hydroxy-2,3,6,8-tetramethoxyxanthone | 24.11 | 333.097 | 92806 |  |  |  |
| **116** | Trigofoenoside A |  |  |  | 24.43 | 901.4799 | 37870 |
| **117** | Sarsasapogenin | 24.71 | 417.3357 | 21534 |  |  |  |
| **118** | Decussatin | 24.89 | 303.0867 | 206980 |  |  |  |
| **119** | α-Linolenic acid | 24.91 | 279.2321 | 2001587 | 24.92 | 277.2168 | 7070 |
| **120** | Vernolic acid | 24.91 | 297.2428 | 941348 | 24.92 | 295.2285 | 132623 |
| **121** | Nimbolide | 25.11 | 467.2073 | 126584 |  |  |  |
| **122** | 7-Hydroxy-5,8-dimethoxyflavone | 25.26 | 299.0916 | 156238 |  |  |  |
| **123** | 9,12,15-Octadecatrienoic acid | 25.27 | 279.2319 | 116111 |  |  |  |
| **124** | Yuccagenin | 25.73 | 431.3152 | 10171 |  |  |  |
| **125** | 6-Desacetylnimbinene | 25.9 | 441.2277 | 282881 |  |  |  |
| **126** | Withanolide B | 26.26 | 455.2779 | 22208 |  |  |  |
| **127** | Momordicoside F2 | 26.5 | 619.4207 | 56662 |  |  |  |
| **128** | Swerchirin | 27.11 | 289.0706 | 117531 |  |  |  |
| **129** | Nimbin | 27.24 | 541.2433 | 158320 |  |  |  |
| **130** | Stearidonic acid | 27.3 | 277.2158 | 139949 | 27.3 | 275.2018 | 12498 |
| **131** | Salannin | 28.57 | 597.3063 | 190292 | 29.09 | 595.2894 | 708292 |
| **132** | (E,E)-9-Oxooctadeca-10,12-dienoic acid | 30.56 | 295.2268 | 165538 | 30.56 | 293.2127 | 136548 |
| **133** | Withanolide G | 31 | 455.2795 | 72709 |  |  |  |
| **134** | Trichosanic acid | 31.12 | 279.2319 | 515764 |  |  |  |
| **135** | Momordicoside K |  |  |  | 34.16 | 647.4157 | 46434 |
| **136** | Momordicine I |  |  |  | 34.89 | 471.3485 | 375438 |
| **137** | γ-Linolenic acid | 36.24 | 279.2321 | 683623 |  |  |  |
| **138** | Betulinic acid |  |  |  | 40.3 | 455.3518 | 34552 |
| **139** | Oleanolic acid | 41.52 | 457.3668 | 231710 | 41.67 | 455.3543 | 173402 |
